# Tango-seq: overlaying transcriptomics on connectomics to identify neurons downstream of *Drosophila* clock neurons

**DOI:** 10.1101/2024.05.22.595372

**Authors:** Alison Ehrlich, Angelina A. Xu, Sofia Luminari, Simon Kidd, Christoph D. Treiber, Jordan Russo, Justin Blau

## Abstract

Knowing how neural circuits change with neuronal plasticity and differ between individuals is important to fully understand behavior. Connectomes are typically assembled using electron microscopy, but this is low throughput and impractical for analyzing plasticity or mutations. Here, we modified the *trans*-Tango genetic circuit-tracing technique to identify neurons synaptically downstream of *Drosophila* s-LNv clock neurons, which show 24hr plasticity rhythms. s-LNv target neurons were labeled specifically in adult flies using a nuclear reporter gene, which facilitated their purification and then single cell sequencing. We call this Tango-seq, and it allows transcriptomic data – and thus cell identity – to be overlayed on top of anatomical data. We found that s-LNvs preferentially make synaptic connections with a subset of the CNMa+ DN1p clock neurons, and that these are likely plastic connections. We also identified synaptic connections between s-LNvs and mushroom body Kenyon cells. Tango-seq should be a useful addition to the connectomics toolkit.

## Introduction

A long-term goal of connectomics is to map a nervous system to help understand how brains function and how variations in brain structure influence individuality and disease.^1^ Starting with simple systems, all of the synapses between the 302 neurons in the *C.elegans* nervous system were reconstructed from EM images.^2^ This reconstruction directly led to insights into behavior – for example refs.^3,4^ and many others. Similarly, the connectome of the *Drosophila* central brain^5^ has deepened our understanding of *Drosophila* brain function, perhaps most notably understanding how innervation subdivides the anatomy of the mushroom body and contributes to learning and memory.^6^

However, electron microscopy has limitations. First, it is still time- and labor-intensive to turn electron microscopy data into a connectome despite recent advances in imaging and analysis. In turn, this means that the available connectomes are a snapshot of a single brain, trapped in time. Thus, EM cannot easily reveal variability between animals or how different states change neuronal connections. In addition, connectomes such as the *Drosophila* hemibrain^5^ often undercount synapses as very fine neurites are difficult to trace. Finally, the connectome only contains chemical synapses and does not include connections mediated by gap-junctions or neuromodulators.

Genetic methods offer higher throughput ways to map neuronal connections. For example, the GRASP method uses two non-fluorescent parts of GFP that are expressed in different neurons. Fluorescence is only detected when two neurons connect and reconstitute GFP across a synapse.^7,8^ Although this method requires deciding on genetically defined sets of candidate neurons in which to express the GFP fragments, GRASP has been used successfully to identify novel connections – for example, downstream of *Drosophila* circadian clock neurons.^9^

*Trans*-Tango is a more recently developed genetic technique for monosynaptic anterograde tracing in *Drosophila*.^10^ In this method, a synaptically-localized human glucagon ligand (hGCG) is expressed in defined neurons using the Gal4/UAS system. hGCG binds a membrane-tethered human glucagon receptor (hGCGR), which is expressed pan-neuronally as a fusion protein with the transcription factor QF. Binding to hGCG to hGCGR-QF recruits a human β-arrestin-TEV protease fusion protein – also expressed pan-neuronally – which cleaves the receptor-QF fusion protein, allowing QF to enter the nucleus and activate a QUAS reporter gene (see Figure 1A). *Trans*-Tango was successfully used to identify second order taste neurons in *Drosophila*,^10^ outputs of the mushroom body,^11^ and neurons involved in light-mediated regulation of locomotor activity.^12^ *trans*-Tango has recently been adapted to map neural circuits in zebrafish.^13^

**Figure 1:**
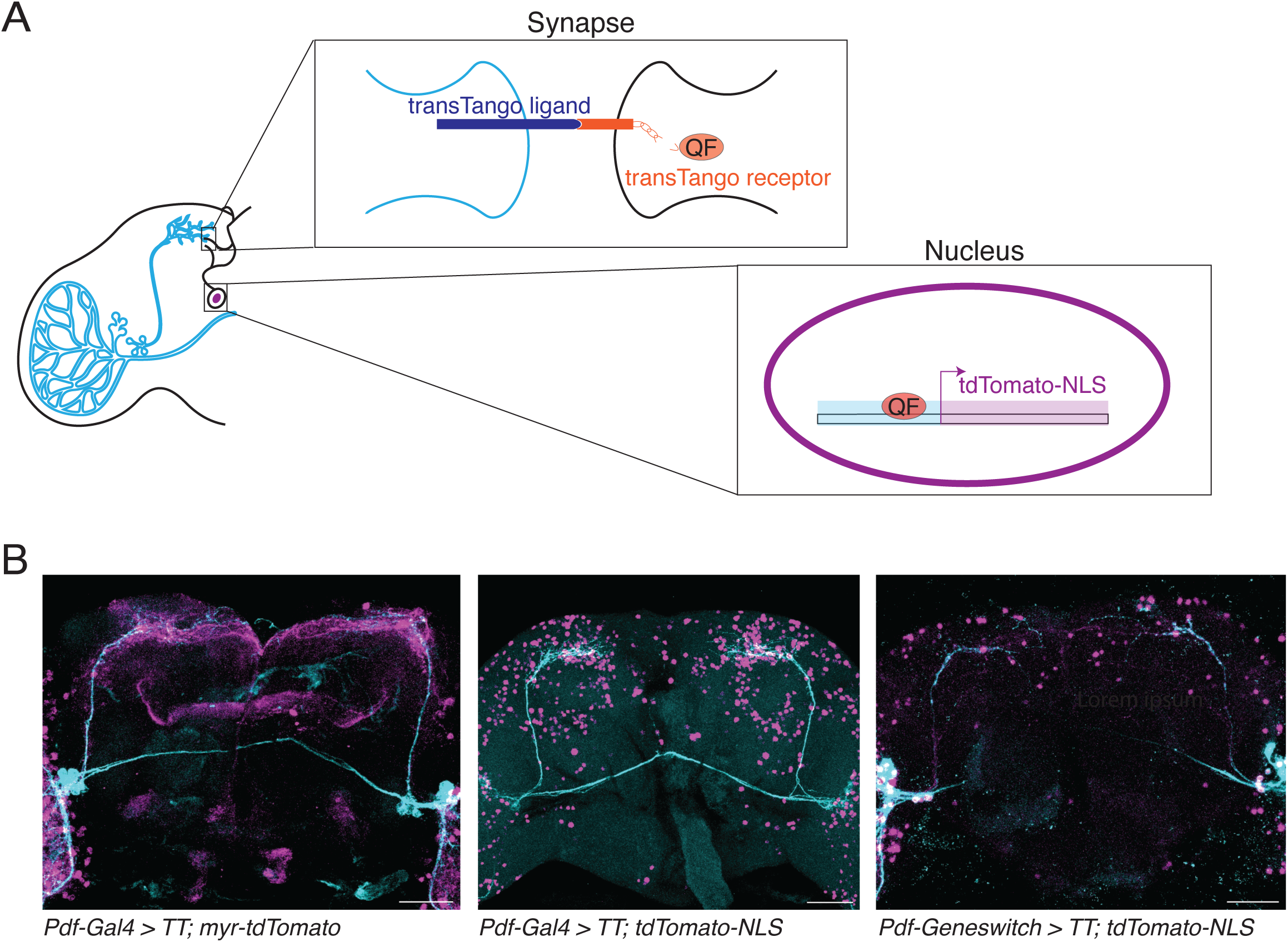
Modifying *trans-*Tango to map s-LNv target neurons. A: Schematic representation of the *trans-*Tango system, with LNvs (light blue, left) expressing the *trans-*Tango ligand. Interaction of the *trans-*Tango ligand and receptor at a synapse with a downstream neuron releases the transcription factor QF (orange, middle), which translocates to the nucleus to activate a reporter gene which is a nuclear-localized tdTomato reporter gene, in this case (right). B: Maximum projection of single midbrains from *trans-Tango* flies with: Left: *Pdf-Gal4* and the original *QUAS-myr-tdTomato* reporter gene Middle: *Pdf-Gal4* and the *QUAS-tdTomato-NLS* reporter gene Right: *Pdf-GS* and the *QUAS-tdTomato-NLS* reporter gene with flies switched to mifepristone-containing food 5-9 days after eclosion. Antibodies to GFP (cyan) label the presynaptic s-LNv and l-LNv neurons. Antibodies to RFP (magenta) label their postsynaptic targets. Brains were dissected and fixed at ZT2. Scale bars = 50 μm.

The small lateral ventral neurons (s-LNvs) are key circadian clock pacemaker neurons in *Drosophila* and are involved in behaviors with 24 hour rhythms including locomotor activity and feeding.^14,15^ s-LNvs show dramatic structural plasticity with their dorsal projections defasciculating and growing in three dimensions at dawn, and then refasciculating and retracting at dusk.^16^ s-LNv plasticity includes making and breaking synaptic connections with a 24 hour rhythm^9^ and the ability to expand s-LNv dorsal projections is required for some circadian behaviors.^12,17^ The s-LNvs express the neuropeptide Pigment Dispersing Factor (PDF) that is critical for circadian rhythms.^14^ s-LNvs also use glycine as a neurotransmitter,^18^ and blocking synaptic transmission can alter adult locomotor activity in certain light conditions.^19^

We decided to use *trans*-Tango to identify s-LNv synaptic targets. We modified the original *trans*-Tango method^10^ by using an inducible Gal4 to ensure that we only detected synaptic connections of s-LNvs in adult flies, and by using a nuclear-localized reporter gene to facilitate cell counting. Our data indicate sparse synaptic connectivity between s-LNvs and other clock neuron groups, which may be a general feature of the circadian clock network. We purified fluorescently-labeled s-LNv target neurons by FACS and used single cell sequencing to obtain their transcriptomes and reveal their identity. We call this method Tango-seq, and it revealed that s-LNvs preferentially synaptically connect with the CNMa-expressing subset of DN1p clock neurons. We also found an interesting synaptic connection between s-LNvs and the Kenyon cells of the mushroom body. Finally, we show that the modified nuclear trans-Tango assay can be used to identify s-LNv synaptic connections that change with structural plasticity.

## Results and Discussion

### Optimizing *trans*-Tango to map adult synaptic connections

We first used the *Pdf-Gal4* driver, which is expressed in only eight neurons per hemisphere in the mature adult central brain: four small and four large LNvs.^20^ The data in Figure 1B using the original membrane-bound *trans-*Tango fluorescent reporter gene (*QUAS-myr-tdTomato*^10^) shows quite broad Tomato reporter gene expression in the adult brain. To more clearly visualize these neurons downstream of the LNvs, we generated a QUAS-tdTomato reporter gene which is localized to the nucleus. A similar nuclear QUAS reporter was recently used to examine second order taste neurons in *Drosophila*.^21^ We refer to *trans-*Tango experiments using the QUAS-tdTomato-NLS reporter as “n*trans*-Tango”. *Pdf-Gal4* expression of n*trans*-Tango still revealed a large number of neurons expressing Tomato, which was surprising given the recent description of 55 medium-and high-strength connections to s-LNvs in the hemibrain electron microscopy connectome.^22^

In addition to the adult LNvs, *Pdf-Gal4* is expressed in 4 tritocerebral neurons which undergo programmed cell death in the first few days post eclosion.^23,24^ *Pdf-Gal4* is also expressed in the 4 larval LNvs in each hemisphere,^20^ which have functional molecular clocks, contribute to larval light avoidance,^25^ and become the adult s-LNvs.^26^ Therefore, some neurons detected using *Pdf-Gal4* in Figure 1B could be connections that existed before adulthood or during the first days of adult life, but are not maintained in the mature adult brain. To test this idea, we used the GeneSwitch (GS) system^27^ to control when the *trans-*Tango ligand is expressed. The GS fusion protein contains the ligand-binding domain of the Progesterone receptor, and is thus inactive as a transcriptional activator until flies ingest Mifepristone, also known as RU486. Mifepristone is easy to add to fly food at relatively low doses and so can be used to temporally control when a UAS transgene is expressed without other effects to adult flies.^27^

We used *Pdf-GeneSwitch* (*Pdf-GS* ^28^) to induce expression of the trans-Tango ligand in mature adult flies. Flies were typically 5-9 days old when transferred to mifepristone-containing food. The images in Figure 1B from brains with *Pdf-GS* expressing n*trans*-Tango still show many cells in the central brain, but fewer than with *Pdf-Gal4*. We imaged brains taken from *Pdf-GS* expressing n*trans*-Tango with food that contained the EtOH vehicle but not mifepristone, but did not detect any reporter gene expression (Figure S1A). We tested how long flies need access to mifepristone for robust expression of tdTomato. While the upstream UAS reporter was activated after ∼1 day on mifepristone, strong expression of the downstream QUAS reporter required 5-7 days (Figure S1A).

The lower numbers of cells expressing the nTomato reporter when using *Pdf-GS* compared to *Pdf-Gal4* could be at least partly due to LNv connections during development. To test this, we raised *Pdf-GS* > n*trans*-Tango flies on mifepristone-containing food so that they express the *trans*-Tango ligand throughout development. Flies were then dissected in the first 24hr of adult life. The data in Figure S1B show strong Tomato expression in the nuclei of many neurons in the adult brain. Since induction of reporter gene expression in *Pdf-GS* expressing n*trans*-Tango flies takes several days (Figure S1A), the expression pattern seen on day 1 of adult life is most likely due to LNv connections formed during development. We therefore used *Pdf-GS* for the remaining experiments to identify s-LNv targets in the adult brain, with flies aged for 5-9 days and then placed on mifepristone food for at least 7 days.

### s-LNvs sparsely connect with other clock neuron types

We first sought to identify which clock neurons are downstream of s-LNvs. For this, we isolated brains with *Pdf-GS* expressing n*trans*-Tango shortly after dusk at Zeitgeber Time 14 (ZT14, two hours after lights off in a 12:12 Light-Dark cycle). We chose ZT14 so that brains could be co-stained with antibodies to the circadian transcription factor Vrille (Vri), which marks all clock neurons and peaks around dusk.^29^ The data in Figure 2A show myr-GFP labeled LNvs in cyan, the tdTomato-NLS labeled downstream neurons in magenta, and the Vri-expressing clock neurons in yellow. Neurons appear white when Vri and tdTomato are co-expressed. These data show that there are LNv targets in many groups of clock neurons. However, there are also many clock neurons that do not express tdTomato, even when LNvs target some neurons within that clock neuron subset.

**Figure 2:**
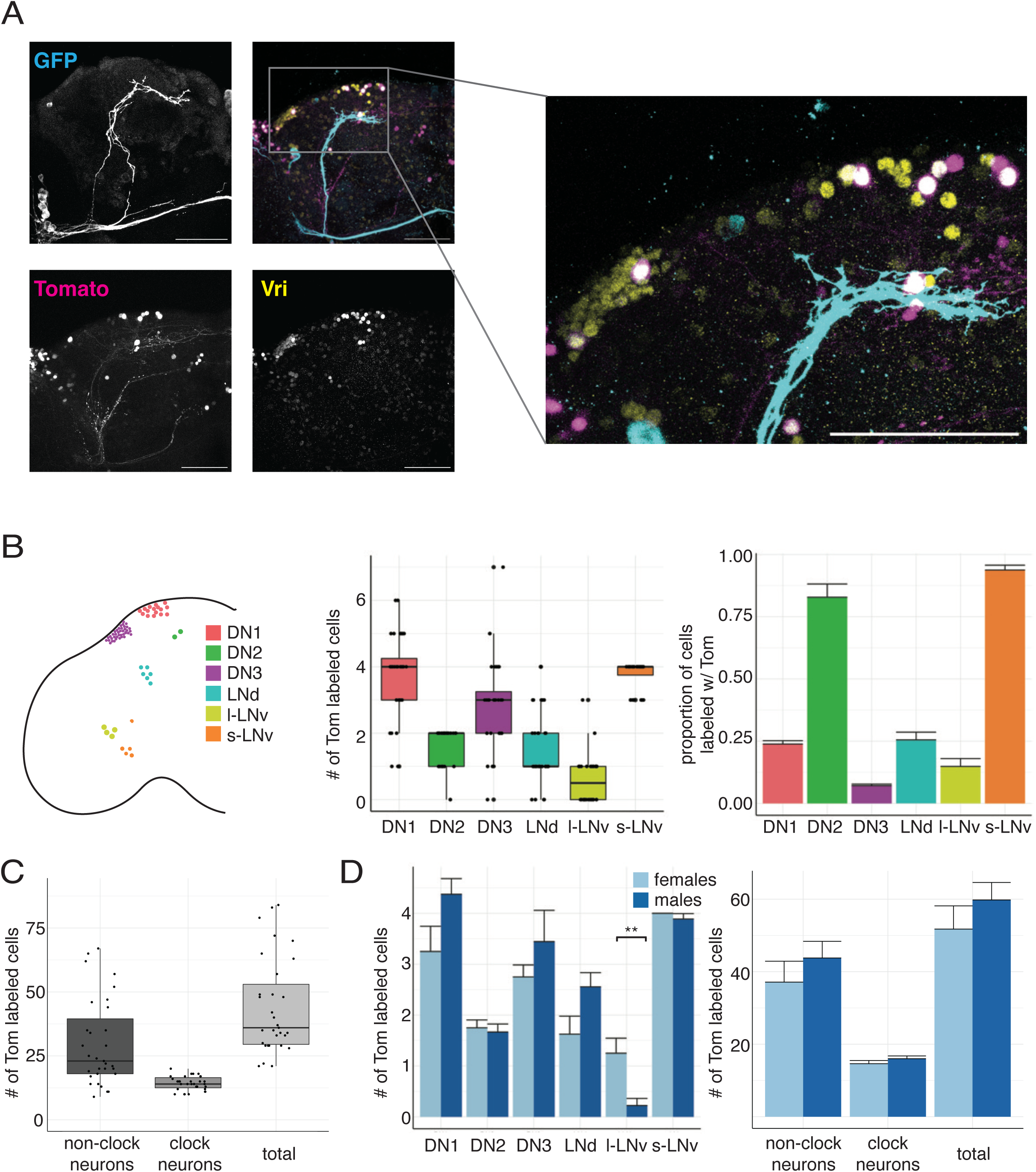
s-LNvs make sparse synaptic connections on to other clock neuron groups. A. Left: Maximum projection of confocal images of brains dissected and fixed at ZT14 from *Pdf-GS > UAS-myr-GFP, trans-Tango; QUAS-tdTomato-NLS* flies with *Pdf-GS* induced for > 7 days. Antibody staining is shown for the GFP+ LNvs (cyan), tdTomato (magenta), and Vri (yellow) channels as well as a merged image. Right: Magnified z-stack of highlighted area showing dorsal clock neuron clusters. Scale bars = 50 μM. B: Color coded schematic of clock neuron types showing approximate location and number of cells for six clock neuron types (left). Graphs quantify the number of tdTomato labeled neurons from each clock cell type (center) and proportion (right) of tdTomato labeled neurons from each clock cell type (N=31 hemispheres). C: Quantification of the number of non-clock neurons, clock neurons, and total neurons labeled with tdTomato (N = 31 hemispheres). D: Quantification of the number (left) and proportion (right) of clock neuron types labeled with tdTomato separated by sex (N = 9 hemispheres for males and N = 8 hemispheres for females). There is a statistically significant difference (Student’s *t*-test, P<0.01) for the number of l-LNvs labeled between male and female flies. The difference is not significant between sexes (P>0.05) for any other cell type. In box and whisker plots, box includes 1^st^ quartile, median, and 3^rd^ quartile values. Whiskers extend to whichever is smaller of +/-1.5 interquartile range or to the max/min value. Bar plots display mean value with error bars indicating standard error of the mean (SEM).

The s-LNvs project to the dorsal protocerebrum of the midbrain, whereas the l-LNvs – which also express *Pdf-GS* – project to the optic lobe.^14^ Thus we only counted neurons in the midbrain to approximate the number of neurons downstream of s-LNvs. The graph in Figure 2B shows the number (left) and the proportion (right) of tdTomato-labeled neurons from each clock neuron type per hemisphere in the midbrain. The data show that s-LNvs are presynaptic to at least six different clock neuron types including themselves. We counted the total number of LNv target neurons in the midbrain and found an average of 14 ± 0.5 clock neurons, 30 ± 3 non-clock neurons, and 44 ± 3 total neurons per hemisphere downstream of LNvs (Figure 2C).

Our estimate for the total number of neurons postsynaptic to the s-LNvs is within the range found by analyzing s-LNv connections in the fly hemibrain connectome.^22^ This study found that individual s-LNvs had at least a medium-strength connection to 55 other neurons. Since each of the 4 s-LNvs has a fairly similar population of target neurons,^22^ our estimate of 44 neurons downstream of s-LNvs per hemisphere is similar to the EM data, and indicates that *Pdf-GS* expression of *ntrans-Tango* likely identifies s-LNv target neurons with strong and medium-strength synaptic connections from s-LNvs.

The data in Figure 2B shows there is a range of connectivity between s-LNvs and different clock neuron groups. At one extreme, we almost always detect all four s-LNvs and both DN2s downstream of s-LNvs. However, other clock neuron groups such as the DN3s show sparse labeling with an average of three of the ∼40 DN3s in each hemisphere labeled with tdTomato. This finding supports the growing idea that there is more heterogeneity than previously appreciated within each anatomical group of clock neurons^30^ – in this case, heterogeneity for s-LNv synaptic connections.

The number of tdTomato-labeled neurons of each type varies between individual brains, which could come from including both male and female brains in the data for Figure 2B. We therefore compared the number of tdTomato-labeled clock neurons in brains isolated separately from female and male flies (Figure 2D). The number of l-LNvs expressing tdTomato differed between female (1.3 ± 0.3 neurons) and male flies (0.2 ± 0.1). However, we cannot distinguish between s-LNvs targeting l-LNvs, or l-LNvs targeting themselves as *Pdf-GS* expresses in both s-and l-LNvs.^28^ There were no significant differences between sexes for any other clock neuron types nor in the total number of neurons, clock neurons, or non-clock neurons between sexes. Therefore, the variability we see in the number of s-LNv targets between brains is unlikely to be caused by sex differences in connectivity. Instead, it is either genuine biological variation between animals, or – perhaps, more likely – due to technical reasons associated with the *trans*-Tango method, GeneSwitch induction, antibody staining and/or imaging.

### DN1p clock neurons also make sparse connections to other clock neurons

We also wanted to test if the sparse synaptic connectivity observed between s-LNvs and other clock neuron types is a general feature of the clock neuron network. To test this, we expressed n*trans-*Tango using a Gal4 line that expresses only in the 6 CNMa+ DN1ps (*CNMa-Gal4*^31^), and a second line expressed in 10-12 DN1ps (*Clk4.1-Gal4*,^32^ which we call *DN1p-Gal4*). The expression patterns of these Gal4 lines are shown in Figure S2A. One caveat of using these Gal4 lines in this experiment is their constitutive expression.

Figure S2B shows brains dissected at ZT14 from these two lines stained for GFP (cyan), tdTomato (magenta), and Vri (yellow). We counted the number and types of clock neurons downstream of these two DN1p subgroups. The data in Figure S2C show that both the smaller CNMa+ population and the larger population marked by *DN1p-Gal4* contact downstream clock neurons in a similar sparse manner to that observed for s-LNvs in Figure 2B. The DN1ps appear to contact both of the DN2s. However, they only contact a small subset of other subtypes – for example, less than 1/5^th^ of the DN3 group of clock neurons.

The cell types synaptically targeted by DN1ps and s-LNvs differ substantially. For example, the s-LNvs synaptically connect to several DN1ps, whereas the DN1ps do not appear to reciprocally connect to s-LNvs – at least not synaptically. Our data suggest that sparse synaptic connections between clock neuron types and their target clock neuron populations is a general feature of the *Drosophila* clock circuit. However, additional Gal4 drivers that cleanly mark other clock neuron subgroups will be required to investigate this idea more fully.

### Tango-seq reveals the molecular identity of cell types downstream of s-LNvs

*Trans-*Tango can identify the anatomical location of downstream neurons, but identifying the cells is difficult without additional markers such as Vri. We decided to take a transcriptomic-based approach to identify s-LNv target neurons. We dissected brains of *Drosophila* with *Pdf-GS* expressing n*trans*-Tango at ZT14, and removed and discarded the optic lobes to enrich for cells in the midbrain, where the s-LNvs project. Brains were dissociated into a single cell suspension and sorted via fluorescent activated cell-sorting (FACS) to isolate tdTomato-labeled neurons. These cells were then processed for single cell RNA sequencing (scRNAseq). We refer to this combination of *trans-*Tango and single scRNAseq as Tango-seq.

Given the small number of neurons per brain labeled by *Pdf-GS* driving n*trans*-Tango, we determined that minimizing false positives was important. Gating for fluorescent cells was stringent and we likely discarded some fluorescent cells as a result. In two biological replicates, we isolated ∼300 and ∼550 cells respectively from approximately 200 individual midbrains in total. With ∼45 labeled cells per midbrain, our sequencing capture rate is less than 10% of the labeled cells. There are many steps between dissection and the final count of sequenced cells, and we likely lose cells at each step.

We performed multiple quality control steps to ensure that all cells included in downstream analysis exceeded a certain sequencing quality. These steps included eliminating cells with fewer than 400 or more than 2500 unique genes (features) expressed. Extremely low feature counts indicate ambient RNA or poor sequencing quality, while high feature counts can indicate a doublet as opposed to a single cell. We also excluded cells whose counts include > 7% mitochondrial gene expression, which increases as cells die and thus can indicate unhealthy cells.^33^ Figure S3 shows the basic quality measures of each dataset before and after trimming. Each biological replicate was independently normalized before variable features were identified.

We used the Canonical Correlation Analysis (CCA) machine learning technique to integrate the two replicates into a single dataset. CCA finds a shared gene correlation structure between two or more datasets by finding linear combinations of features across datasets that are maximally correlated.^34^ We rescaled the data and performed dimensional reduction via principal component analysis (PCA) and uniform manifold approximation and projection (UMAP). Figure S3B shows two dimensions of our UMAP dimensional reduction of the integrated dataset separated by replicate. The salt-and-pepper patterning of cells from each replicate indicates that the two replicates are very similar to each other. Subsequent analyses were performed on the integrated dataset, shown as one dataset in Figure 3A.

**Figure 3:**
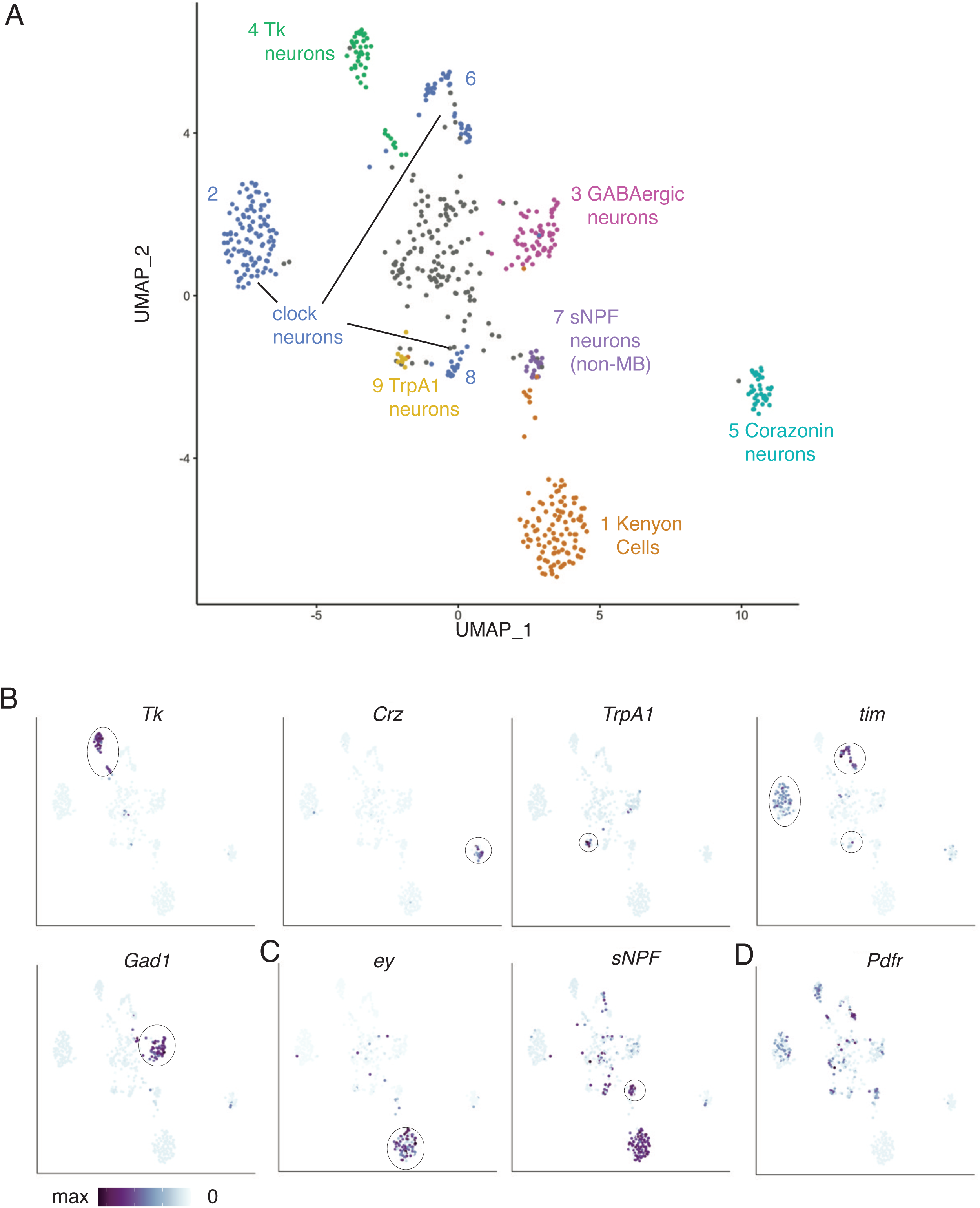
Tango-seq reveals the molecular identity of neurons downstream of s-LNvs. A: UMAP plot indicating single cell sequencing of clusters of tdTomato neurons isolated from *Pdf-GS > UAS-myr-GFP, trans-Tango; QUAS-tdTomato-NLS* brains. *Pdf-GS* was induced with mifepristone for 7 days and brains dissected at ZT14. Optic lobes were discarded during dissection to enrich for s-LNv targets. Transcriptional profiles of each cluster were compared to existing single-cell sequencing atlases.^35,36^ Cell-type information is indicated for 9 of the 10 clusters. B: UMAP plots of normalized gene expression for features identifying the labeled clusters. Color scales range from maximum expression of the indicated gene (purple) to undetectable (white). Most clusters had unique identifying marker genes: *Tk* mRNA identifies Cluster 4; *Crz* identifies Cluster 5; *TrpA1* identifies Cluster 9; *tim* identifies Clusters 2, 6 and 8 as clock neurons; and *Gad1* identifies Cluster 3. C: UMAP plot of normalized expression of *ey* (left) and *sNPF* (right). Co-expression of *ey* and *sNPF* identify Cluster 1 as Kenyon cells of the mushroom body. Neurons with *sNPF* but not *ey* are likely an unrelated population of *sNPF-*expressing neurons. D: UMAP plot showing normalized *Pdf receptor* expression with some clusters not expressing detectable levels of *Pdfr*.

Clustering of the scRNAseq data revealed 10 clusters of cells representing different cell types (Figure 4A). 8 of the 10 clusters have molecular markers that correspond with known cell types. Of the two remaining clusters, cluster 0 – the grey cells in the center of Figure 3A – does not have markers that clearly identify it as a specific cell type and probably comprises cells that are rare targets, cells with relatively low-quality sequencing, or cells erroneously included in the dataset due to imperfect FACS. Cluster 3, the other unidentified cluster, consists of a population of GABAergic neurons that cannot be unambiguously identified as a particular cell type but appear to be the only GABAergic cell type synaptically downstream of s-LNvs.

**Figure 4:**
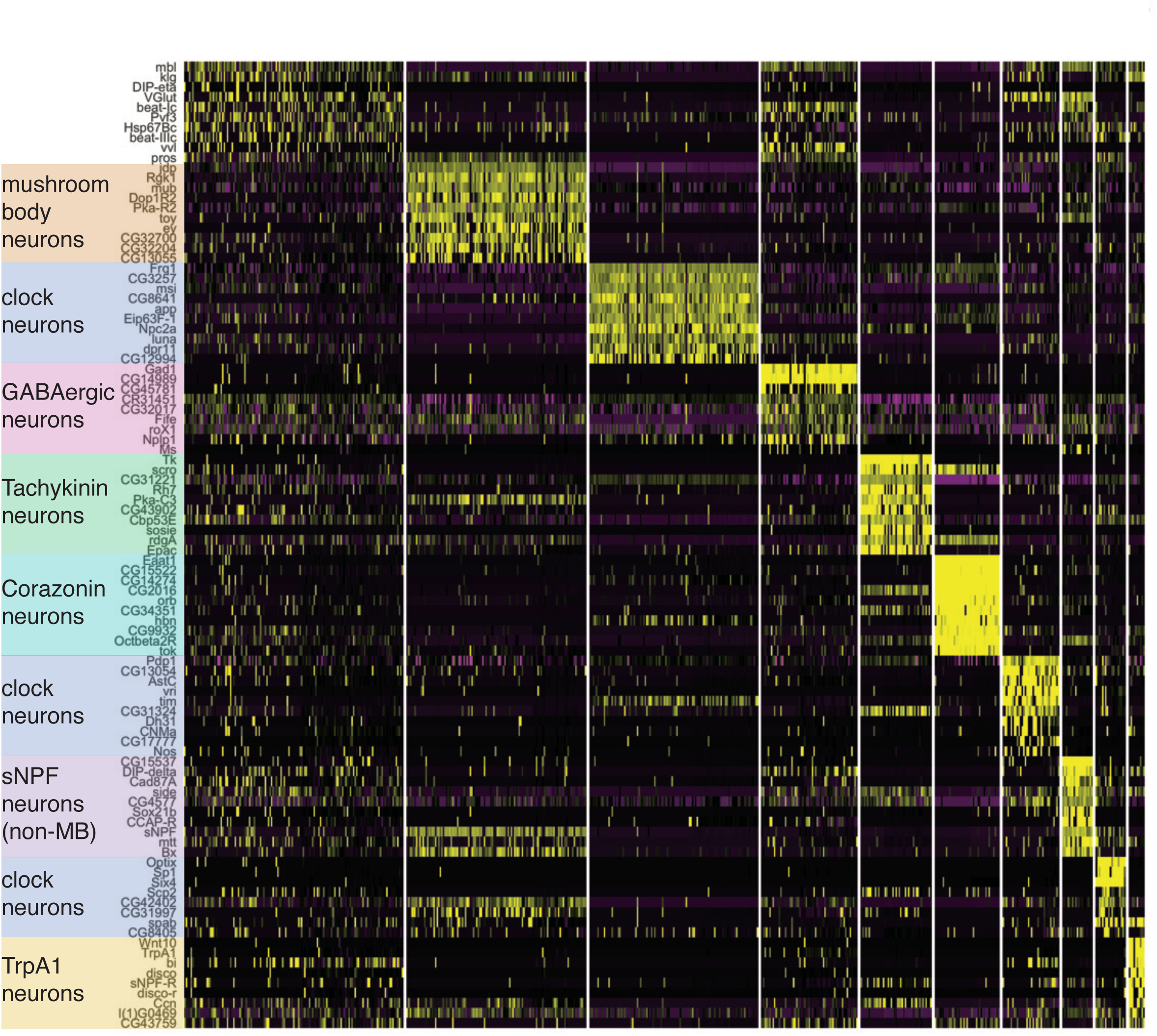
Specific marker genes identify molecular identity of clusters. Heat map of the 10 most upregulated genes per cluster sorted by log2 Fold Change. Identified cell types are indicated (left) and correspond with the labeled clusters in Figure 3 and the more diffuse Cluster 0. Yellow indicates high expression, purple indicates low expression.

Marker genes that identify the 8 identified cell types are shown in Figure 3B-C. Neuropeptides are often expressed in relatively small numbers of neurons in the fly brain and their mRNAs were useful in identifying four different groups of s-LNv target neurons. *Corazonin* (*Crz*) and *Tachykinin* (*Tk*) are neuropeptides that label clusters 3 and 4 in Figure 3. *short neuropeptide-F* (*sNPF*) is more broadly expressed in the fly brain, but its co-expression with *eyeless* (*ey*) in mushroom body (MB) Kenyon cells (KC) identifies cluster 1 as KCs (Figure 3C). Furthermore, expression of genes like *ab*, *GstD11* and *CG13055* indicate that most (>80%) of these KCs are γ-lobe KCs.^35^ Cells in cluster 7 which express *sNPF* but not *ey* are a population of non-MB *sNPF* neurons. Temperature sensitive neurons in the midbrain express the temperature sensitive cation channel *TrpA1* and are present in cluster 9. Clusters 2, 6, and 8 express the clock neuron markers *timeless* (*tim*) and *vri*, identifying these clusters as clock neurons. The top 10 most highly enriched genes for each cluster are shown in Figure 4. We note that the *Crz*-expressing cells in cluster 3 have the same markers as T1 optic lobe neurons.^36^ This is surprising given that s-LNvs project to the dorsal brain and not the optic lobe. However, there are ∼800 T1 neurons in each optic lobe^37^ and we manually dissected the midbrain away from the optic lobe in our experiments. Therefore, it seems likely that our dissections were not perfect and that T1 optic lobe neurons are downstream of l-LNvs rather than s-LNvs.

The neuropeptide PDF is the main signal released from s-LNvs and is required for normal circadian behavior.^14,38^ Consistent with this, the feature plot in Figure 3D shows that *Pdf Receptor* (*Pdfr*) is broadly expressed in s-LNv target neurons. However, *Pdfr* expression was not detected in KCs or T1 neurons, indicating that they likely receive information from LNvs via a different neuropeptide or neurotransmitter.

We did not find all of the cells that we expected using Tango-seq. For example, we did not find the s-LNvs themselves even though they communicate with one another^22,39^ and are labeled with *Pdf-GS* expressing n*trans*-Tango (Figure 1B). This is likely because s-LNvs express both GFP and tdTomato and we avoided sorting cells with high GFP levels as they could be auto-fluorescent cells. We also did not recover the lateral-horn leucokinin (LHLK) neurons despite information from LNvs propagating to LHLK neurons.^40^ This indicates that either s-LNvs do not communicate with LHLKs synaptically, or that LHLK neurons are not directly downstream of sLNvs.

### Sub-clustering of clock neurons reveals specific clock cell types as s-LNv targets

We next wanted to identify precisely which clock neurons were in the three clusters of clock neurons identified as s-LNv target neurons by Tango-seq. For this, we selected all neurons that expressed the clock neuron marker *vri* and re-clustered them into 3 clusters (Figure 5A). We also selected clock neurons using the clock gene *tim.* However, *vri* identified a smaller subset of cells all of which could be unambiguously identified via specific markers, and all neurons that expressed *vri* also expressed *tim* (Figure 5B).

**Figure 5:**
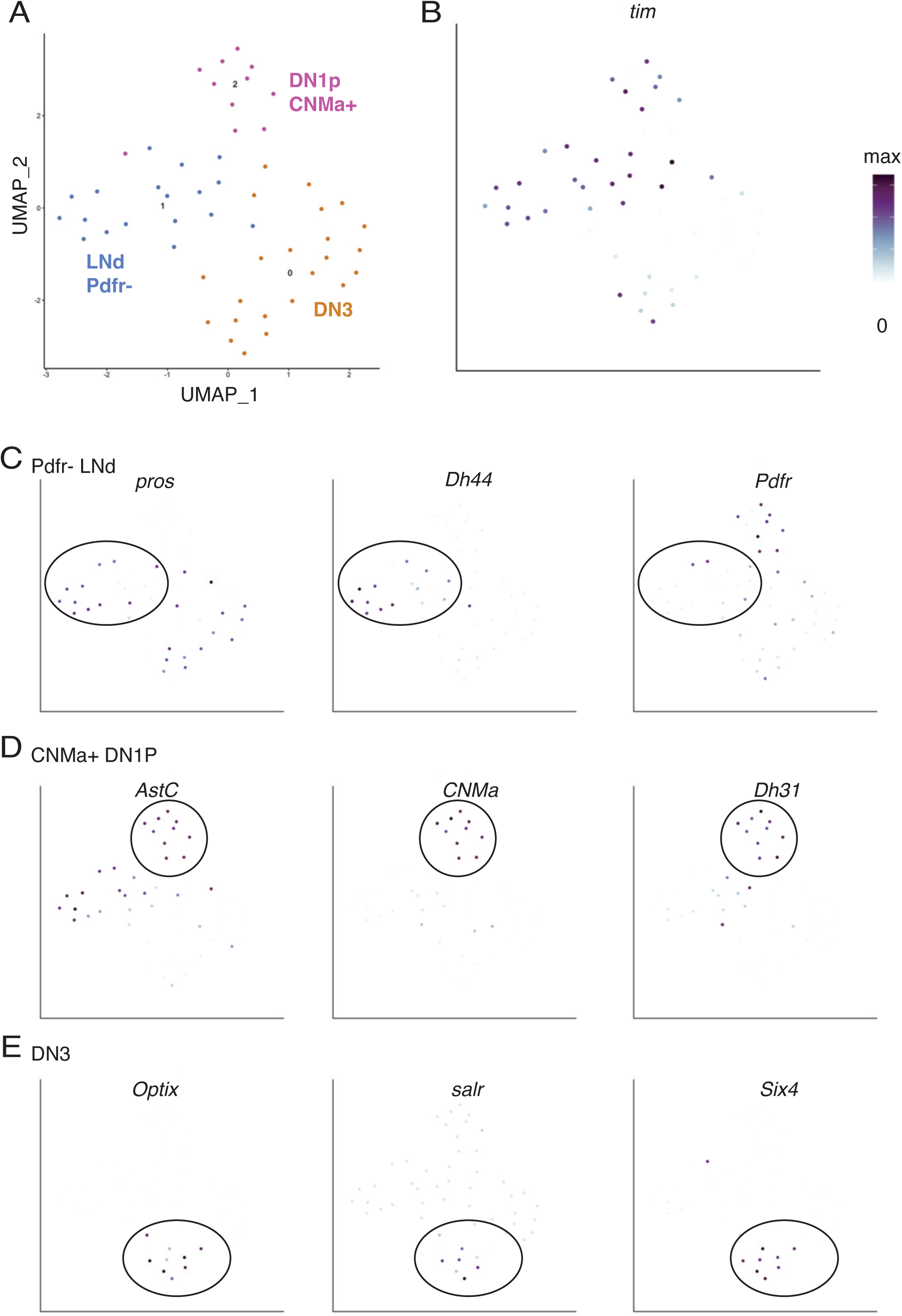
Subclustering of *vrille-*expressing clock neurons reveals details of clock neuron subtypes downstream of s-LNvs. A. All cells with normalized expression of the clock neuron marker *vri* greater than 0.2 were selected from the dataset in Figure 3A. The data was rescaled and clustered. The UMAP plot shows clusters of clock neurons, with three subtypes specifically indicated. B. UMAP shows normalized expression of *tim* in the selected cells. All cells express *tim* at levels above baseline but with variable levels. C. UMAP of positive markers for LNds – *pros* (left) and *Dh44* (center) – as well as for *Pdfr* (right) indicating that most LNds postsynaptic to s-LNvs do not express the PDF receptor. D. UMAP of positive markers for DN1ps including *CNMa* (center) which is only expressed in a subset of DN1p clock neurons.^30^ E. UMAP of *Optix, salr, and Six4* as markers of DN3 clock neurons.^30^

We found marker genes for each of these three clusters and compared these data to an scRNAseq study of all adult clock neurons.^30^ Cluster 1 expresses the transcription factor *prospero* (*pros*) and the neuropeptide *Diuretic hormone 44* (*Dh44*), which marks it as a subgroup of dorsal lateral neurons (LNds) (Figure 5C). Interestingly, most of these cells do not express *Pdfr* at high levels. Since there are three *Pdfr+* and three *Pdfr-* LNds per hemisphere^41,42^, our data indicate that s-LNvs preferentially make strong synaptic connections with the *Pdfr-* subset. This indicates that communication between s-LNvs and these LNds is likely mediated through a signal other than PDF. However, the lack of synaptic connections with *Pdfr+* LNds does not necessarily mean that s-LNvs do not communicate with *Pdfr+* LNds because bath-applied PDF rapidly increases intracellular cAMP in half of the LNds in a *Pdfr*-dependent manner.^43^ Therefore, this communication likely occurs through non-synaptic PDF-signaling.

We identified cluster 2 as the posterior dorsal clock neurons (DN1ps) as they express the neuropeptides *Allatostatin C* (*Ast-C*) and *Diuretic hormone 31* (*Dh31*) (Figure 5D).^30,44^ These cells also express the gene that encodes the neuropeptide CNMamide (*CNMa*).^45^ As we did not find any other DN1ps in our dataset, we conclude that the s-LNvs preferentially make synaptic connections with CNMa+ DN1ps. Finally, we identified cluster 0 as DN3s since they express *Optix*, *spalt-related (salr)*, and *Six4* that are DN3 marker genes^30^ (Figure 5E).

### Validating s-LNv connections identified by Tango-seq

We next wanted to confirm some of the cell types determined by Tango-seq as s-LNv targets. Some connections such as the s-LNv to DN2 connection have been previously described using GRASP^46^ and EM. ^47^ In addition, synaptic connections between s-LNvs and DN1ps are not surprising because of previous data using GRASP.^9,12,48,49^ However, we wanted to test the idea that s-LNvs preferentially connect to the CNMa+ DN1ps. Therefore, we co-stained brains with *Pdf-GeneSwitch* driving n*trans-Tango* with an antibody to CNMa. The data in Figure 6A show that tdTomato is detected strongly in 1 DN1p and more weakly in 2-3 additional DN1ps, all of which produce CNMa (labeled in yellow). We did not detect any obvious tdTomato-expressing DN1ps that do not produce CNMa in this or other brains. These data confirm the idea that s-LNvs preferentially synaptically connect with the CNMa+ DN1ps, although seemingly with only a subset of this subset.

**Figure 6:**
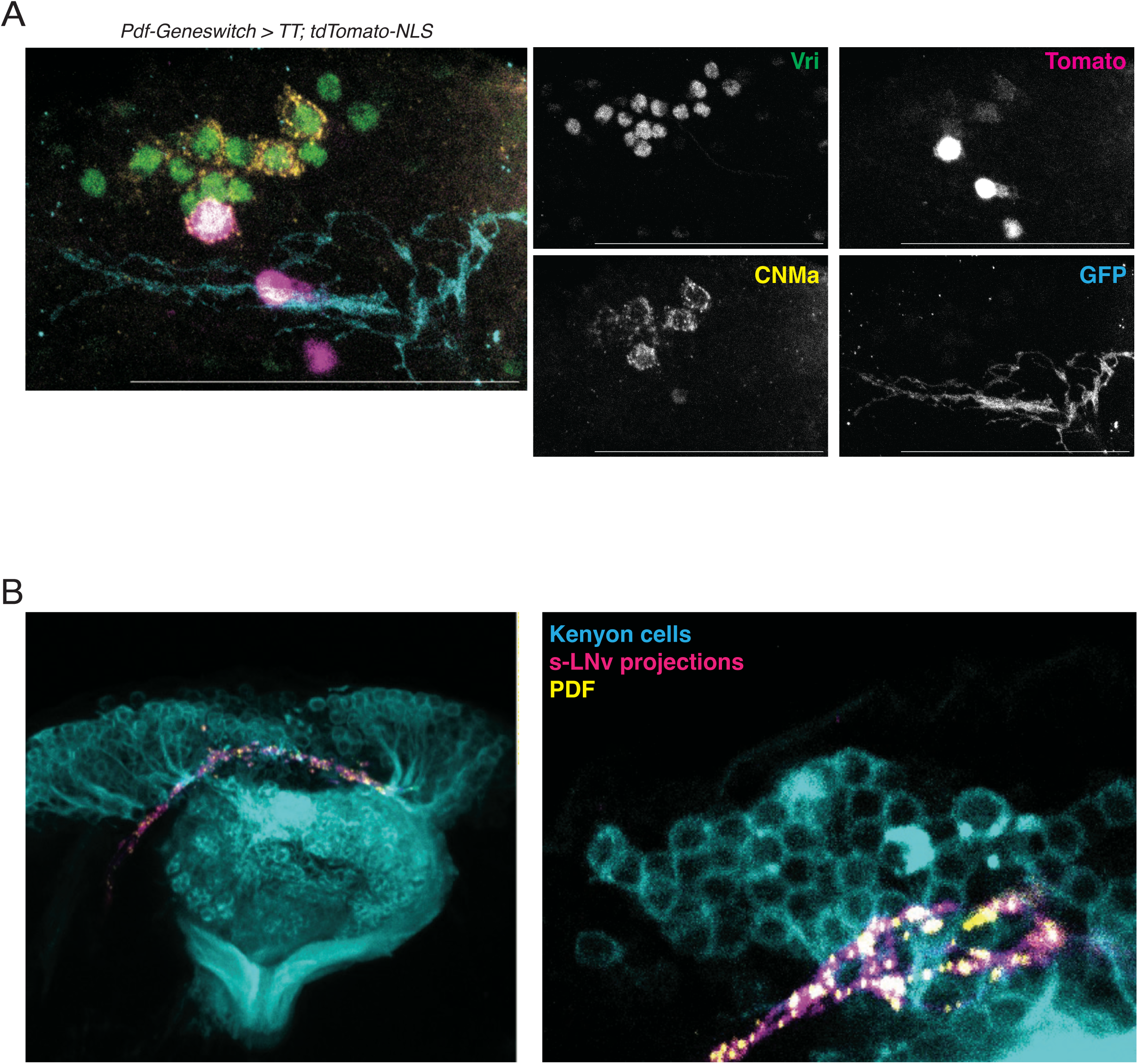
Validating s-LNv target neurons. A: Maximum projection of *Pdf-GS > UAS-myr-GFP, trans-Tango; QUAS-tdTomato-NLS* fly brains at ZT14. Antibodies recognize GFP (cyan) to mark LNv projections, tdTomato (magenta) to mark s-LNv target neurons, Vri (green) to mark clock neurons, and CNMa (yellow) to mark the CNMa-expressing DN1ps. Scale bar = 50 μM. B: Confocal images of s-LNv projections labeled via a *Pdf-RFP* transgene (magenta) and PDF (yellow), and Kenyon cell bodies and dendrites (cyan) labeled via *MB247-Gal4* expressing *UAS-CD8::GFP.* The image on the right shows a maximum projection. The image on the right is from a second brain and shows a 3μm z-stack with s-LNv projections in the same plane as Kenyon cells.

The mushroom body the main learning and memory center in *Drosophila*.^50^ There are circadian rhythms in learning and memory^51,52^, and circadian rhythms of gene expression in the mushroom body that depend on the clock gene *period*, which is not expressed by Kenyon cells.^53^ In addition, a GRASP signal was found between s-LNvs and two different Gal4 drivers that express in the mushroom body.^9^ Thus, it is perhaps not surprising that Tango-seq detected direct synaptic connections between s-LNvs and the mushroom body KCs (Figure 3-4). However, the *Drosophila* hemibrain connectome did not report any direct s-LNv connections to KCs.^5^

To test if s-LNvs project to KCs independently of *trans*-Tango, we used flies with a *Pdf-RFP* transgene to label s-LNvs and their projections, and the membrane-targeted *CD8-GFP* expressed via *MB247-Gal4* to label Kenyon cell bodies and their projections into the mushroom body. We also added antibody to the PDF neuropeptide since PDF puncta often overlap with the pre-synaptic marker Bruchpilot.^17,54^ Figure 6B shows a confocal image of the s-LNv projections in magenta and the KCs and their projections in cyan. The s-LNv projections pass through the cell-body layer of the MB, very close to the KCs (Figure 6B). These data are consistent with the Tango-Seq data showing that s-LNvs synaptically connect with KCs (Figure 3-4). The data are also consistent with our exploration of the *Drosophila* hemibrain connectome, which revealed that the s-LNv projections pass over the MB calyx and into the cell body layer (Figure S4). However, this was not a brain area analyzed for synaptic connections.^5^

### Using n*trans*-Tango to detect plasticity-induced changes

The low throughput nature of electron microscopy makes it challenging to test how connections between neurons change during structural plasticity. We therefore wanted to test if n*trans-* Tango could be used to test how s-LNv connectivity changes during s-LNv plasticity. The projections of s-LNvs are expanded at dawn and retracted at dusk.^16^ However, the n*trans-* Tango reporter gene takes several days to accumulate to sufficient levels to be detected (Figure S1A), making it unrealistic to visualize differences in s-LNv connectivity between dawn and dusk.

Instead, we decided to use n*trans-*Tango to measure synaptic connections when s-LNv projections are constitutively retracted and compare the resulting pattern to control s-LNvs which expand and retract. s-LNv projections are constitutively retracted in flies with either a null mutation in the clock gene *period* (*per*)^16^ or via adult-specific RNAi knockdown of *per* or *timeless* (*tim*).^55^ We raised flies as normal to ensure the clock circuit develops normally, and then moved flies to constant light (LL) for 3 days before inducing expression of the trans-Tango ligand using *Pdf-GS* and mifepristone for a further 7 days in LL. Constant light prevents accumulation of the light-sensitive Tim protein which, in turn, prevents accumulation of Per, which requires Tim for its stability and nuclear entry.^56^

To test if the s-LNv projections in flies exposed in LL are in a contracted state, we measured the 3D spread of s-LNv projections at ZT2 (dawn), ZT14 (dusk) and compared them to flies kept in LL for 10 days. s-LNv projections were quantified using the membrane-bound GFP reporter gene co-expressed with the trans-Tango ligand. The data in Figure 7A show that s-LNvs expressing nTrans-tango show normal structural plasticity rhythms in LD, with their projections more expanded at ZT2 (dawn) than at ZT14 (dusk).^16^ We found that the 3D spread of s-LNv projections of flies in LL is different from flies isolated at ZT2, but not different from flies isolated at ZT14. Therefore, we conclude that LL keeps s-LNv retractions in a dusk-like retracted state.

**Figure 7:**
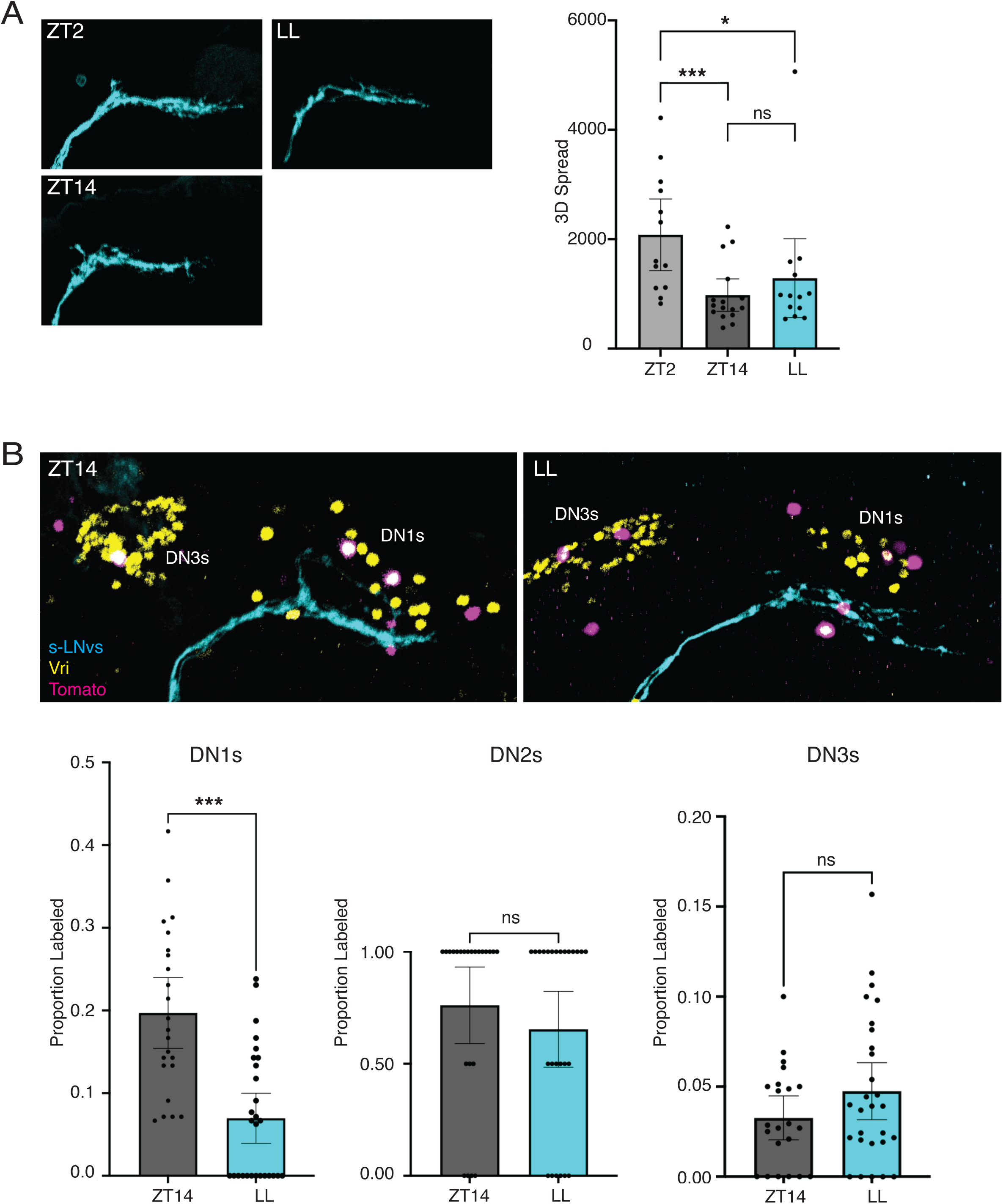
n*t*rans-Tango identifies s-LNv target neurons whose connections are altered by s-LNv plasticity. A: Left: Representative s-LNv projections from brains of *Pdf-GS > UAS-myr-GFP, trans-Tango; QUAS-tdTomato-NLS* from flies isolated at ZT2 (left top), ZT14 (left bottom) or after 10 days in constant light (LL, right). Right: Quantification of the 3D spread of s-LNv projections in the graph on the right shows that s-LNv projections at ZT2 are more expanded than at ZT14 or in LL. The data show quantification of at least 13 hemispheres from 2 independent experiments. B: Top: s-LNv target neurons imaged from *Pdf-GS > UAS-myr-GFP, trans-Tango; QUAS-tdTomato-NLS* flies isolated at ZT14 (left) or after 10 days in constant light (LL, right) with the last 7 days on mifepristone-containing food. s-LNv projections are marked via GFP (cyan), downstream neurons via Tomato (magenta) and clock neurons via Vri (yellow). Bottom: Quantification of the proportion of DN1s (left), DN2s (middle) and DN3s (right) that are s-LNv targets when s-LNv projections are allowed to expand and retract (ZT14, grey) or when constitutively retracted (LL, blue). The data show quantification of at least 24 hemispheres from 3 independent experiments.

Next, we compared the number of TdTomato-expressing clock neurons in flies with *Pdf-GS* expressing n*trans*-Tango at ZT14 with flies maintained in LL as above. Although s-LNv projections are retracted at ZT14, the TdTomato signal is stable and the clock marker Vri is easily detectable at ZT14 and in LL, as it is in *per* and *tim* mutant flies.^29^ The presence of Vri allowed us to count the proportion of visible clock neurons co-labelled with tdTomato. This is important because it is sometimes not possible to image all DN1ps because of the orientation of the brain. Therefore, we wanted to ensure that any differences between brains isolated from flies at ZT14 and LL are due to changes in synaptic connectivity rather than not being able to visualize all DN1s.

The data in Figure 7B show an average of 3.3 Tomato-expressing DN1s in brains isolated at ZT14, compared to only 1.1 Tomato-expressing DN1s in flies kept in LL. There was no significant difference between the numbers of Tomato-expressing DN2s or DN3s between ZT14 and LL. Thus, we conclude that keeping s-LNv projections in a constitutively retracted state reduces connections between s-LNvs and DN1ps. Given that s-LNvs preferentially connect to the CNMa+ DN1ps, the sLNv to CNMa+ DN1p connection is likely one of the plastic connections normally made and broken during the daily structural plasticity of s-LNvs. There may also be differences in s-LNv connectivity with non-clock neurons between ZT14 and LL, but this will require identifying additional markers to label and count these cells.

### Conclusions and unanswered questions

The combination of anatomical data and single cell sequencing that we report here supports previously published data on s-LNv connectivity such as the s-LNv to DN2^46^ and s-LNv to DN1p connections.^9,12,48,49^ It also provides new insights into s-LNv synaptic targets, such as preferential connections to CNMa+ DN1ps, and connections to KCs. Our study also raises many questions including the nature of the code that helps s-LNvs connect to a specific subset of DN1ps – especially if this is a plastic connection made and broken every day. Mechanistic questions that address the molecular basis of neuronal connectivity should be answerable using n*trans*-Tango. In addition, the behavioral functions of many of the s-LNv synaptic connections that we describe here are also unknown. For example, does the s-LNv to γ-KC connection explain circadian rhythms in memory formation and recall?^51,52^ Overall, we believe Tango-seq is a useful addition to the genetic toolkit for dissecting neural circuits, especially in combination with adult-restricted expression of the *trans*-Tango ligand.

## Materials and methods

### Fly strains and rearing

All *Drosophila melanogaster* lines were raised at 25°C in 12:12 light:dark cycles unless otherwise specified. Although strongest expression of the *trans*-Tango reporter gene was previously reported for flies reared at 18°C,^10^ we raised flies at 25°C because some neurons form more synapses and have more synaptic partners when raised at 18°C.^57^ Fly genotypes and their sources are listed in Table 1. To induce *Pdf-GS* flies were moved to food containing mifepristone dissolved in ethanol to a final concentration of 500 μg/mL. Flies were typically kept on food containing mifepristone for 7-14 days.

**Table 1:** Fly lines.

| Genotype | Source |
| --- | --- |
| *Pdf*-GS / CyO | BDSC stock #81116 |
| *Pdf*-Gal4 | Jae Park<sup>20</sup> |
| *CNMa*-Gal4 | Greg Suh<sup>31</sup> |
| *Sco* / CyO; *Clk4.1*-Gal4 | BDSC stock #36316 |
| *UAS*-mGFP, *QUAS*-mtdTomato; *trans*-Tango; | BDSC stock #77124 |
| *trans*-Tango; *UAS*-mGFP, *QUAS*-tdTomato-NLS | This study |
| *MB247*-Gal4, *UAS*-CD8::GFP | This study |
| *Pdf*-RFP | Ref. <sup>58</sup> |

### Generating transgenic lines

The UAS-myr-GFP, QUAS-tdTomato-NLS plasmid was cloned using Infusion HD Cloning Kit (Takara) and sent to Genetivision Corporation for injection. Reporter transgenes containing a mini *white^+^* selection marker were inserted into the attP2 site on 3L using PhiC31 recombinase mediated methods.^59^ The tdTomato coding sequence is codon-optimized for *Drosophila.*^60^

### Immunohistochemistry, confocal imaging and quantification

Fly brains were dissected at specified times of day in cold phosphate-buffered saline (PBS) and fixed and permeabilized using standard procedures.^17^ Brains were incubated with primary antibodies (see Table 2 below) overnight at 4°C, and with secondary antibodies either overnight at 4°C or at room temperature for 1-2 hours. Brains were mounted in SlowFade Gold antifade reagent (Invitrogen). Confocal images were acquired using a Leica SP8 confocal microscope. Images were taken at 1 μm slices using a 20x or 63x oil immersion lens. z-stacks were produced in FIJI by combining the maximum intensity for each pixel in each channel. Cell counting was performed manually using the FIJI cell counter plugin. Clock cell types were identified by staining with antibodies to Vri and their position in the brain. The 3D spread of s-LNv projections in Figure 7 used a custom MATLAB script.^17^

**Table 2:** Primary Antibodies.

| Target | Host | Concentration | Source/ identifier |
| --- | --- | --- | --- |
| PDF | Mouse | 1:50 | AB_2315084 |
| Vri | Guinea Pig | 1:5000 | Paul Hardin |
| GFP | Sheep | 1:500 | NovusBio NB100-62622 |
| GFP | Chicken | 1:500 | Aveslab #GFP-1020 |
| RFP | Rabbit | 1:500 | MBL #PM005 |
| RFP | Mouse | 1:100 | MBL #M204-7 |
| CNMa | Rabbit | 1:500 | Greg Suh |

### Sequencing

#### Sample preparation

Brains were dissected from male and female flies in PBS. Approximately 100 brains were dissected by at least three individual dissectors over the course of an hour. Dissected brains were kept on ice in Schneider’s insect media. The optic lobes were removed and discarded. Brains were dissociated into a single-cell suspension by incubating in 2 mg/mL collagenase and 2 mg/mL dispase in PBS for 1.5 hours at 25 °C in a shaking dry-block incubator. The enzymes were then carefully removed, and brains were washed twice with 400 μL of ice-cold Schneider’s insect media (Sigma-Aldrich). Schneider’s media was removed and replaced with ice cold PBS + 0.04% BSA. The brains were pipetted up and down until there were no large chunks of tissue remaining. The cells were then filtered through a 20-µm cell strainer. Balancing the need to maximize manual dissociation without harming the cells and the need for filtration to achieve a single cell suspension were likely substantial sources of cell dropout. Cells were kept on ice between each step. DAPI was added to the cell suspension to a final concentration of 1 μg/mL. Cells were sorted on the BDS FACSAria II at NYU’s CGSB GenCore. Single cells were selected by forward scatter gating. Dead cells were excluded by not selecting cells that had taken up DAPI, which was typically ∼30% of cells. tdTomato+ cells were selected by stringent gating for RFP.

#### Library preparation and sequencing

Preparation of single cell transcriptomes was performed using the Chromium Next GEM Single Cell 3ʹ Reagent Kits v3.1 according to the protocol described in the Chromium Next GEM Single Cell 3ʹ v3.1(Dual Index) User Guide • Rev A. Due to the small number of tdTomato labeled cells per brain, we expected a final cell recovery of <1000 cells per experiment. We produced two independent libraries as biological replicates. Library quantification and quality control were performed using high sensitivity D1000 ScreenTape and reagents on the Agilent TapeStation 4200 to examine the fragment size distribution for the initial cDNA library amplification and for the final sequencing library. Additional size selection bead cleanups (SPRIselect, Beckman Coulter) were performed as necessary to remove any primer dimers that could interfere with sequencing.

Libraries were prepared for dual end sequencing using the i7 and i5 primers from Dual Index TT Set A plate and were run on the Illumina NextSeq 500 using a NextSeq 500/550 Mid Output Kit v2.5 (150 Cycles) with the recommended number of cycles from the 10x Genomics user guide (28 x 10 x 10 x 90). Cells were sequenced to an average depth of greater than 180,000 reads per cell.

#### Bioinformatic analyses

Sequencing libraries were demultiplexed and processed using the Cell Ranger-6.0.1 pipeline and aligned to the *Drosophila melanogaster* dm6 transcriptome. The resultant single cell count matrices were analyzed using Seurat V4 ^61,62^. Initial quality control on the cell count matrices included eliminating cells with <400 or >2500 genes (features) per cell. These low or high feature counts can indicate a cell fragment or doublet respectively. We also eliminated all cells with >7% of counts from mitochondrial genes, as increased mitochondrial gene expression is associated with cell death. Raw data and our analysis code is available on request.

## Acknowledgements

We are indebted to Orie Shafer for suggesting that we use *Pdf-GS*, and to Ishmail Abdus-Saboor for suggesting the constant light experiment. We are very grateful to Shane Liddelow and Michael Rosbash for numerous insightful comments on the work. We thank Robin Hiesinger for presenting unpublished work on the effect of temperature on synaptic connectivity in an online seminar, Gilad Barnea for developing the trans-Tango system and for sending us flies, and Scott Waddell for his comments and for generously supporting CDT. We thank Fernanda Ceriani, Paul Hardin, Sebastian Kadener, Jae Park, Greg Suh and the Bloomington *Drosophila* Stock Center for flies, DNA and antibodies. This investigation was conducted in facilities constructed with support from the NIH National Center for Research Resources. AE was partly supported by NINDS predoctoral fellowship T32 NS086750. AX, SL and JR were all supported by NYU CAS Dean’s Undergraduate Research Fellowships. CDT was supported by funds from an ERC Advanced Grant (789274) and a Wellcome Collaborative Award (209235) to Scott Waddell (Oxford, UK). This work was supported by NIH grant GM136363 to JB.

---

**Figure S1:**
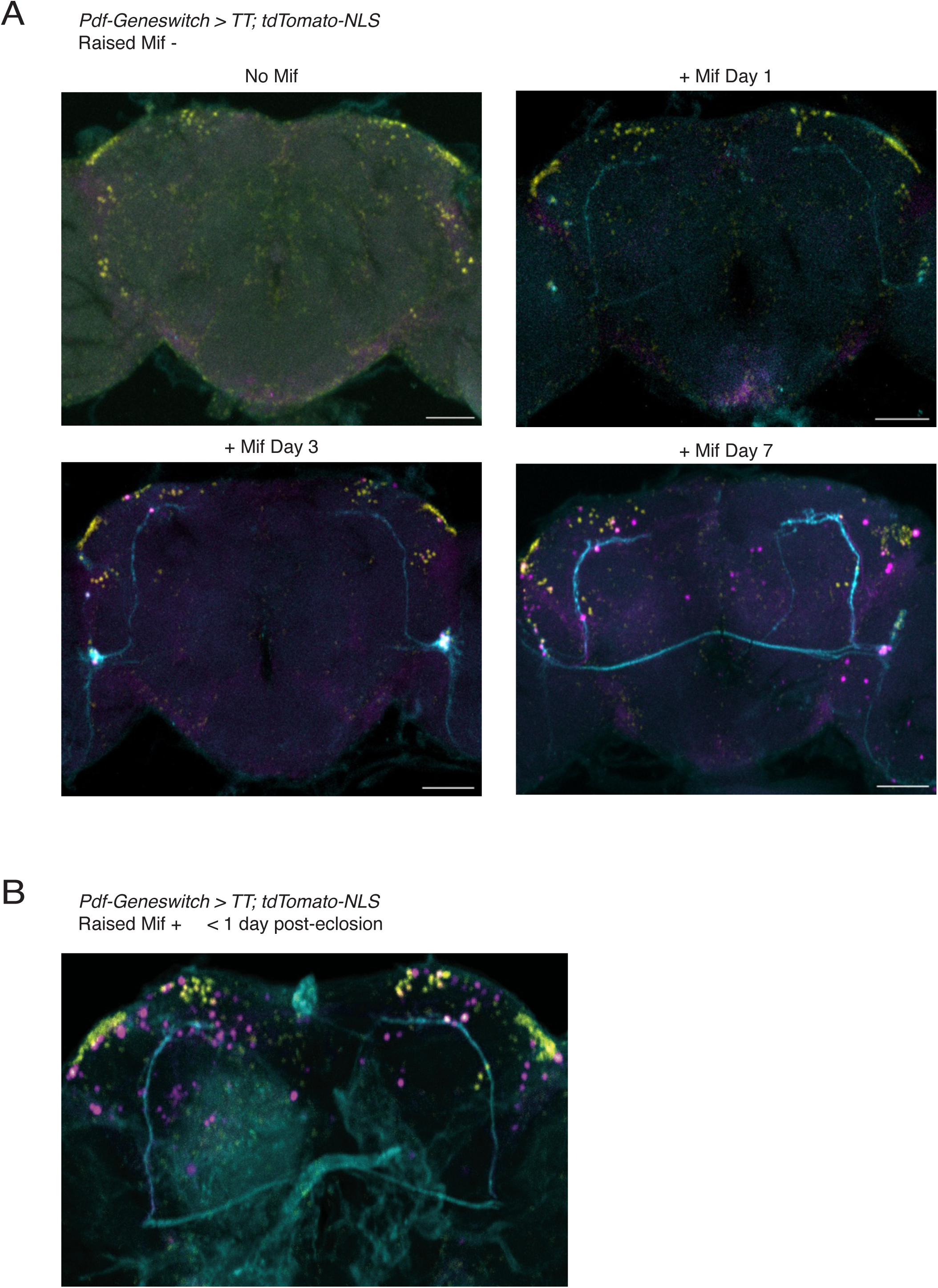
Timecourse of expression of the postsynaptic nuclear tdTomato reporter with *Pdf-GS* used to express the *trans*-Tango ligand. A: *Pdf-GS > trans-Tango; UAS-mGFP, QUAS-tdTomato-NLS* flies were raised on standard food and then switched to food with 500 μg/mL mifepristone added and dissected after a 1, 3 or 7 days, or switched to fly food with 500 μg/mL EtOH (vehicle) for the no mifepristone sample. Antibodies to GFP (cyan) label the LNvs and antibodies to RFP (magenta) label postsynaptic targets. Antibodies to Vri (yellow) label the nuclei of clock neurons. Brains were dissected and fixed at ZT14 when Vri levels are high. B: *Pdf-GS > trans-Tango; UAS-mGFP, QUAS-tdTomato-NLS* flies were raised on food with 500 μg/mL mifepristone so that *Pdf-GS* is active throughout development. Adult flies were dissected at ZT14 on the first day after hatching from the pupal case, and stained as above.

**Figure S2:**
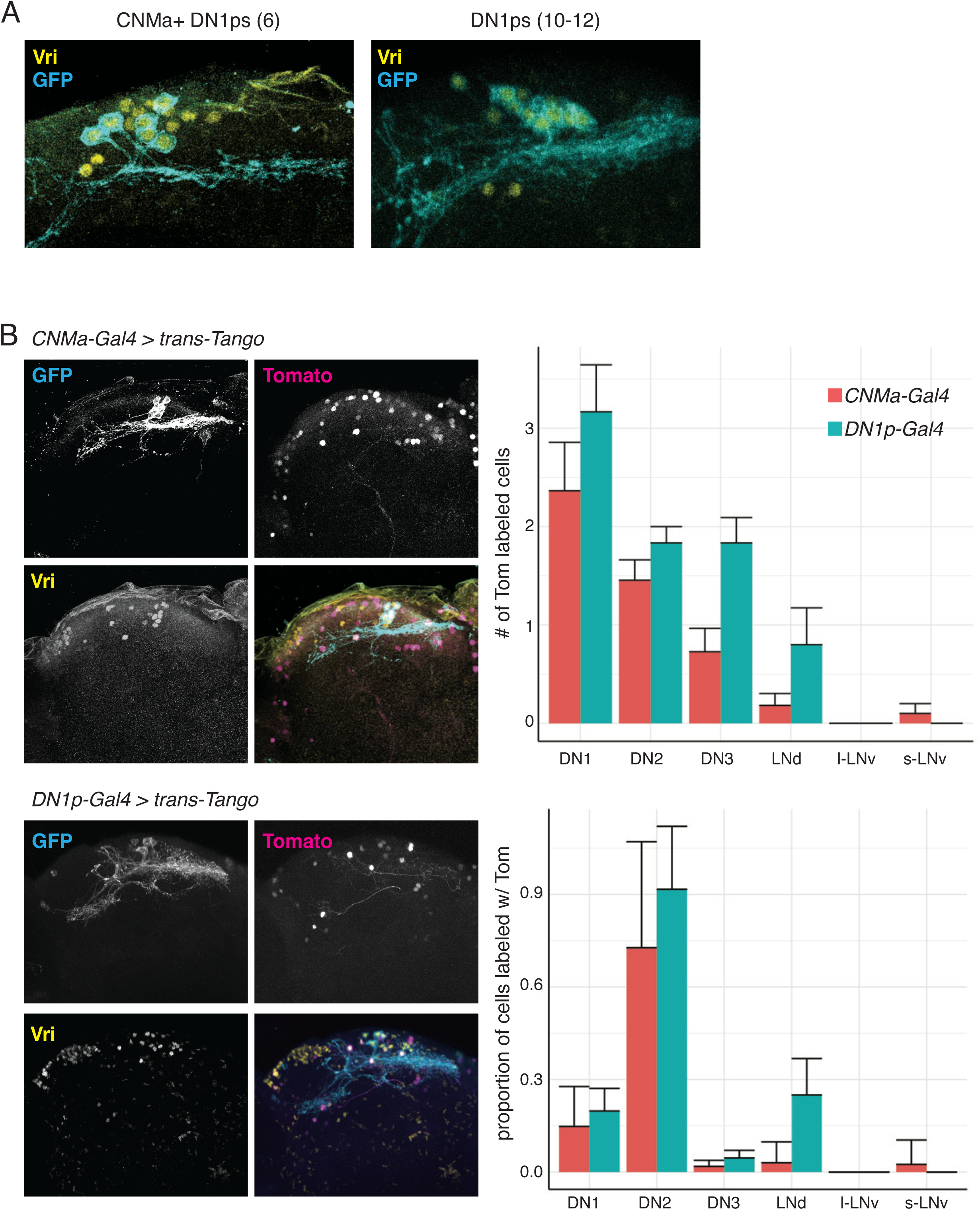
*trans*-Tango using two different DN1p lines shows sparse synaptic connectivity to other clock neuron groups. A: Maximum projections of *CNMa-Gal4* > *UAS-myr-GFP* (left) and *DN1-Gal4 > UAS-myr-GFP* (right). GFP (cyan) and Vri (yellow) are shown for each line. The *CNMa-Gal4* labels 6 neurons per hemisphere and the *DN1p-Gal4* labels 10-12 DN1ps per hemisphere. B. Confocal z-stack of fly brains from *DN1p-Gal4 > UAS-myr-GFP, trans-Tango; QUAS-tdTomato-NLS* (top) *and CNMa-Gal4 > UAS-myr-GFP, trans-Tango; QUAS-tdTomato-NLS* (bottom). Representative antibody staining for GFP (cyan), tdTomato (magenta), and Vri (yellow) are shown as well as merged images (right). C. Graphs quantify the number (top) and proportion (bottom) of tdTomato-labeled neurons from each clock cell type (N=10 hemispheres for *CNMa-Gal4* and N=5 hemispheres for *DN1p-Gal4*). Bar plots display mean value with error bars indicating SEM.

**Figure S3:**
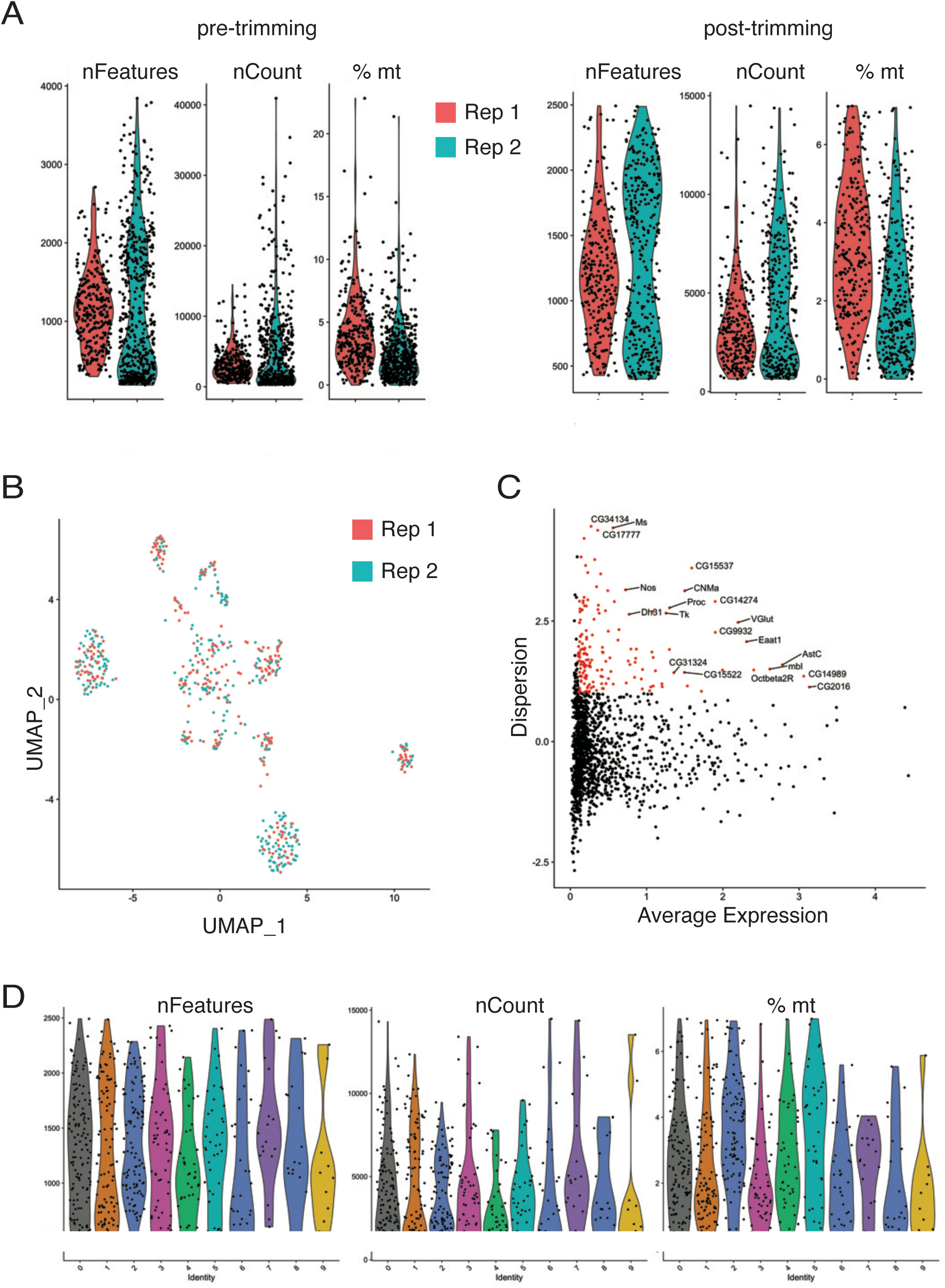
Biological sequencing replicates integrate well and produce a high-quality dataset for post-integration analysis. A: Graphs show violin plots of the number of individual genes identified (nFeatures), number of individual sequencing reads mapped to a transcript (nCount), and the percentage of reads from mitochondrial genes (% mt) before (left) and after trimming (right) the dataset for each cell separated for biological replicates 1 (red) and 2 (blue). B: Uniform manifold approximation and projection (UMAP) plot showing Canonical correlation analysis (CCA) based integration of the two biological replicates with individual cells color-coded by replicate as in A. C: Volcano plot shows genes with high variability between cells in the integrated dataset. Significant variability is a factor of overall expression level of a gene and a z-score derived from the dispersion (a function of variance). The top 20 most variable genes are labeled. D: Violin plots showing the nFeatures, nCount, and % mt for individual clusters derived from the integrated data set. The consistency in these features between clusters indicate they are unlikely to be driving cluster assignment.

**Figure S4:**
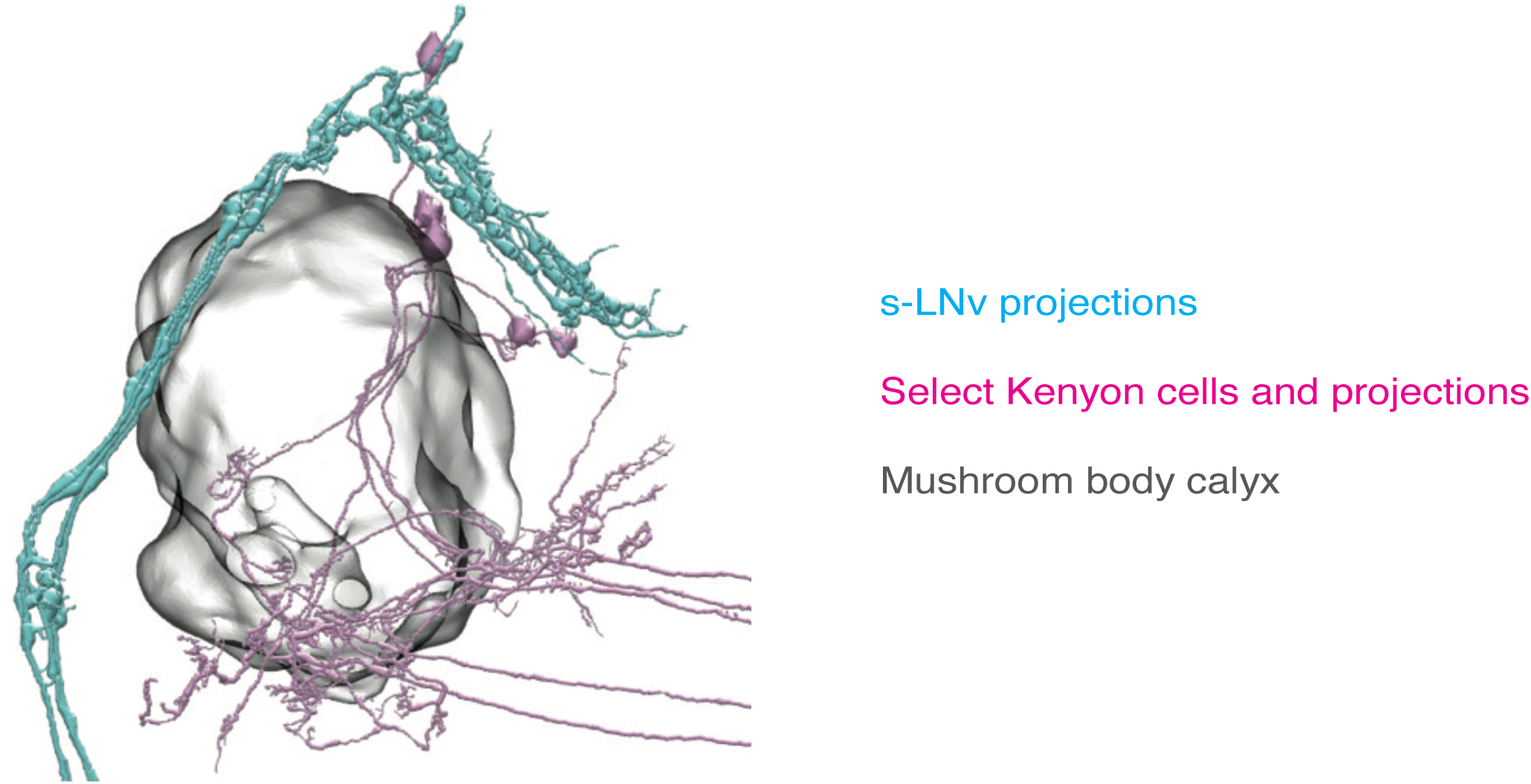
s-LNv projections are very close to γ lobe Kenyon cells. A reconstruction from the hemibrain (v1.2.1) viewer showing the 4 s-LNvs from the right hemisphere (blue), the outline of the right mushroom body calyx (grey), and a selection of γ lobe Kenyon cells (pink). The s-LNv projections pass over the calyx and into the cell body layer of the MB, close to γ lobe Kenyon cells.

## Notes

### Competing Interest Statement

The authors have declared no competing interest.

## References

1. Sporns, O., Tononi, G., and Kotter, R. (2005). The human connectome: A structural description of the human brain. PLoS Comput Biol 1, e42. 10.1371/journal.pcbi.0010042.

2. White, J.G., Southgate, E., Thomson, J.N., and Brenner, S. (1986). The structure of the nervous system of the nematode *Caenorhabditis elegans*. Philos Trans R Soc Lond B Biol Sci 314, 1–340. 10.1098/rstb.1986.0056.

3. Mori, I., and Ohshima, Y. (1995). Neural regulation of thermotaxis in *Caenorhabditis elegans*. Nature 376, 344–348. 10.1038/376344a0.

4. Gray, J.M., Hill, J.J., and Bargmann, C.I. (2005). A circuit for navigation in *Caenorhabditis elegans*. PNAS 102, 3184–3191. 10.1073/pnas.0409009101.

5. Scheffer, L.K., Xu, C.S., Januszewski, M., Lu, Z., Takemura, S.Y., Hayworth, K.J., Huang, G.B., Shinomiya, K., Maitlin-Shepard, J., Berg, S., et al. (2020). A connectome and analysis of the adult *Drosophila* central brain. eLife 9. 10.7554/eLife.57443.

6. Li, F., Lindsey, J.W., Marin, E.C., Otto, N., Dreher, M., Dempsey, G., Stark, I., Bates, A.S., Pleijzier, M.W., Schlegel, P., et al. (2020). The connectome of the adult *Drosophila* mushroom body provides insights into function. eLife 9. 10.7554/eLife.62576.

7. Feinberg, E.H., Vanhoven, M.K., Bendesky, A., Wang, G., Fetter, R.D., Shen, K., and Bargmann, C.I. (2008). GFP Reconstitution Across Synaptic Partners (GRASP) defines cell contacts and synapses in living nervous systems. Neuron 57, 353–363. 10.1016/j.neuron.2007.11.030.

8. Shearin, H.K., Quinn, C.D., Mackin, R.D., Macdonald, I.S., and Stowers, R.S. (2018). t-GRASP, a targeted GRASP for assessing neuronal connectivity. J Neurosci Methods 306, 94–102. 10.1016/j.jneumeth.2018.05.014.

9. Gorostiza, E.A., Depetris-Chauvin, A., Frenkel, L., Pirez, N., and Ceriani, M.F. (2014). Circadian pacemaker neurons change synaptic contacts across the day. Current Biology: 24, 2161–2167. 10.1016/j.cub.2014.07.063.

10. Talay, M., Richman, E.B., Snell, N.J., Hartmann, G.G., Fisher, J.D., Sorkac, A., Santoyo, J.F., Chou-Freed, C., Nair, N., Johnson, M., et al. (2017). Transsynaptic mapping of second-order taste neurons in flies by trans-Tango. Neuron 96, 783–795 e784. 10.1016/j.neuron.2017.10.011.

11. Scaplen, K.M., Talay, M., Fisher, J.D., Cohn, R., Sorkac, A., Aso, Y., Barnea, G., and Kaun, K.R. (2021). Transsynaptic mapping of *Drosophila* mushroom body output neurons. eLife 10. 10.7554/eLife.63379.

12. Song, B.J., Sharp, S.J., and Rogulja, D. (2021). Daily rewiring of a neural circuit generates a predictive model of environmental light. Sci Adv 7. 10.1126/sciadv.abe4284.

13. 13. Coomer, C., Naumova, D., Talay, M., Zolyomi, B., Snell, N., Sorkac, A., Chanchu, J.M., Cheng, J., Roman, I., Li, J., et al. (2023). Transsynaptic labeling and transcriptional control of zebrafish neural circuits. bioRxiv. 10.1101/2023.04.03.535421.

14. Renn, S.C., Park, J.H., Rosbash, M., Hall, J.C., and Taghert, P.H. (1999). A *pdf* neuropeptide gene mutation and ablation of PDF neurons each cause severe abnormalities of behavioral circadian rhythms in *Drosophila*. Cell 99, 791–802. 10.1016/s0092-8674(00)81676-1.

15. 15. Barber, A.F., Fong, S.Y., Kolesnik, A., Fetchko, M., and Sehgal, A. (2021). *Drosophila* clock cells use multiple mechanisms to transmit time-of-day signals in the brain. PNAS 118. 10.1073/pnas.2019826118.

16. Fernandez, M.P., Berni, J., and Ceriani, M.F. (2008). Circadian remodeling of neuronal circuits involved in rhythmic behavior. PLoS Biology 6, e69. 10.1371/journal.pbio.0060069.

17. Petsakou, A., Sapsis, T.P., and Blau, J. (2015). Circadian Rhythms in Rho1 activity regulate neuronal plasticity and network hierarchy. Cell 162, 823–835. 10.1016/j.cell.2015.07.010.

18. Frenkel, L., Muraro, N.I., Beltran Gonzalez, A.N., Marcora, M.S., Bernabo, G., Hermann-Luibl, C., Romero, J.I., Helfrich-Forster, C., Castano, E.M., Marino-Busjle, C., et al. (2017). Organization of circadian behavior relies on glycinergic transmission. Cell Reports 19, 72–85. 10.1016/j.celrep.2017.03.034.

19. Umezaki, Y., Yasuyama, K., Nakagoshi, H., and Tomioka, K. (2011). Blocking synaptic transmission with tetanus toxin light chain reveals modes of neurotransmission in the PDF-positive circadian clock neurons of *Drosophila melanogaster*. J Insect Physiol 57, 1290–1299. 10.1016/j.jinsphys.2011.06.004.

20. Park, J.H., Helfrich-Forster, C., Lee, G., Liu, L., Rosbash, M., and Hall, J.C. (2000). Differential regulation of circadian pacemaker output by separate clock genes in *Drosophila*. PNAS 97, 3608–3613. 10.1073/pnas.97.7.3608.

21. Snell, N.J., Fisher, J.D., Hartmann, G.G., Zolyomi, B., Talay, M., and Barnea, G. (2022). Complex representation of taste quality by second-order gustatory neurons in *Drosophila*. Current Biology: 32, 3758–3772 e3754. 10.1016/j.cub.2022.07.048.

22. Shafer, O.T., Gutierrez, G.J., Li, K., Mildenhall, A., Spira, D., Marty, J., Lazar, A.A., and Fernandez, M.P. (2022). Connectomic analysis of the *Drosophila* lateral neuron clock cells reveals the synaptic basis of functional pacemaker classes. eLife 11. 10.7554/eLife.79139.

23. Helfrich-Forster, C. (1997). Development of Pigment-dispersing hormone-immunoreactive neurons in the nervous system of *Drosophila melanogaster*. J. Comparative Neurology 380, 335–354. 10.1002/(sici)1096-9861(19970414)380:3<335::aid-cne4>3.0.co;2-3.

24. Gatto, C.L., and Broadie, K. (2011). Fragile X Mental Retardation Protein is required for programmed cell death and clearance of developmentally-transient peptidergic neurons. Developmental biology 356, 291–307. 10.1016/j.ydbio.2011.05.001.

25. Collins, B., Kane, E.A., Reeves, D.C., Akabas, M.H., and Blau, J. (2012). Balance of activity between LNvs and glutamatergic dorsal clock neurons promotes robust circadian rhythms in *Drosophila*. Neuron 74, 706–718. 10.1016/j.neuron.2012.02.034.

26. Kaneko, M., and Hall, J.C. (2000). Neuroanatomy of cells expressing clock genes in *Drosophila*: transgenic manipulation of the period and timeless genes to mark the perikarya of circadian pacemaker neurons and their projections. The Journal of comparative neurology 422, 66–94. 10.1002/(sici)1096-9861(20000619)422:1<66::aid-cne5>3.0.co;2-2.

27. Osterwalder, T., Yoon, K.S., White, B.H., and Keshishian, H. (2001). A conditional tissue-specific transgene expression system using inducible GAL4. PNAS 98, 12596–12601. 10.1073/pnas.221303298.

28. Depetris-Chauvin, A., Berni, J., Aranovich, E.J., Muraro, N.I., Beckwith, E.J., and Ceriani, M.F. (2011). Adult-specific electrical silencing of pacemaker neurons uncouples molecular clock from circadian outputs. Current Biology: 21, 1783–1793. 10.1016/j.cub.2011.09.027.

29. Cyran, S.A., Buchsbaum, A.M., Reddy, K.L., Lin, M.C., Glossop, N.R., Hardin, P.E., Young, M.W., Storti, R.V., and Blau, J. (2003). *vrille*, *Pdp1*, and *dClock* form a second feedback loop in the *Drosophila* circadian clock. Cell 112, 329–341. 10.1016/s0092-8674(03)00074-6.

30. Ma, D., Przybylski, D., Abruzzi, K.C., Schlichting, M., Li, Q., Long, X., and Rosbash, M. (2021). A transcriptomic taxonomy of *Drosophila* circadian neurons around the clock. eLife 10. 10.7554/eLife.63056.

31. Kim, B., Kanai, M.I., Oh, Y., Kyung, M., Kim, E.K., Jang, I.H., Lee, J.H., Kim, S.G., Suh, G.S.B., and Lee, W.J. (2021). Response of the microbiome-gut-brain axis in *Drosophila* to amino acid deficit. Nature 593, 570–574. 10.1038/s41586-021-03522-2.

32. Zhang, Y., Liu, Y., Bilodeau-Wentworth, D., Hardin, P.E., and Emery, P. (2010). Light and temperature control the contribution of specific DN1 neurons to *Drosophila* circadian behavior. Current Biology: 20, 600–605. 10.1016/j.cub.2010.02.044.

33. Ilicic, T., Kim, J.K., Kolodziejczyk, A.A., Bagger, F.O., McCarthy, D.J., Marioni, J.C., and Teichmann, S.A. (2016). Classification of low quality cells from single-cell RNA-seq data. Genome Biol 17, 29. 10.1186/s13059-016-0888-1.

34. Butler, A., Hoffman, P., Smibert, P., Papalexi, E., and Satija, R. (2018). Integrating single-cell transcriptomic data across different conditions, technologies, and species. Nature Biotechnology 36, 411–420. 10.1038/nbt.4096.

35. Croset, V., Treiber, C.D., and Waddell, S. (2018). Cellular diversity in the *Drosophila* midbrain revealed by single-cell transcriptomics. eLife 7. 10.7554/eLife.34550.

36. Konstantinides, N., Kapuralin, K., Fadil, C., Barboza, L., Satija, R., and Desplan, C. (2018). Phenotypic convergence: distinct transcription factors regulate common terminal features. Cell 174, 622–635 e613. 10.1016/j.cell.2018.05.021.

37. Naidu, V.G., Zhang, Y., Lowe, S., Ray, A., Zhu, H., and Li, X. (2020). Temporal progression of *Drosophila* medulla neuroblasts generates the transcription factor combination to control T1 neuron morphogenesis. Developmental Biology 464, 35–44. 10.1016/j.ydbio.2020.05.005.

38. 38. Mertens, I., Vandingenen, A., Johnson, E.C., Shafer, O.T., Li, W., Trigg, J.S., De Loof, A., Schoofs, L., and Taghert, P.H. (2005). PDF receptor signaling in *Drosophila* contributes to both circadian and geotactic behaviors. Neuron 48, 213–219. 10.1016/j.neuron.2005.09.009.

39. Collins, B., Kaplan, H.S., Cavey, M., Lelito, K.R., Bahle, A.H., Zhu, Z., Macara, A.M., Roman, G., Shafer, O.T., and Blau, J. (2014). Differentially timed extracellular signals synchronize pacemaker neuron clocks. PLOS Biology 12, e1001959. 10.1371/journal.pbio.1001959.

40. Cavey, M., Collins, B., Bertet, C., and Blau, J. (2016). Circadian rhythms in neuronal activity propagate through output circuits. Nature Neuroscience 19, 587–595. 10.1038/nn.4263.

41. Yoshii, T., Todo, T., Wulbeck, C., Stanewsky, R., and Helfrich-Forster, C. (2008). Cryptochrome is present in the compound eyes and a subset of *Drosophila*’s clock neurons. J. Comparative Neurology 508, 952–966. 10.1002/cne.21702.

42. Im, S.H., Li, W., and Taghert, P.H. (2011). PDFR and CRY signaling converge in a subset of clock neurons to modulate the amplitude and phase of circadian behavior in *Drosophila*. PloS One 6, e18974. 10.1371/journal.pone.0018974.

43. Yao, Z., and Shafer, O.T. (2014). The *Drosophila* circadian clock is a variably coupled network of multiple peptidergic units. Science 343, 1516–1520. 10.1126/science.1251285.

44. Lamaze, A., and Stanewsky, R. (2019). DN1p or the “fluffy” cerberus of clock outputs. Front Physiol 10, 1540. 10.3389/fphys.2019.01540.

45. Jin, X., Tian, Y., Zhang, Z.C., Gu, P., Liu, C., and Han, J. (2021). A subset of DN1p neurons integrates thermosensory inputs to promote wakefulness via CNMa signaling. Current Biology: 31, 2075–2087 e2076. 10.1016/j.cub.2021.02.048.

46. Tang, X., Roessingh, S., Hayley, S.E., Chu, M.L., Tanaka, N.K., Wolfgang, W., Song, S., Stanewsky, R., and Hamada, F.N. (2017). The role of PDF neurons in setting the preferred temperature before dawn in *Drosophila*. eLife 6. 10.7554/eLife.23206.

47. Reinhard, N., Schubert, F.K., Bertolini, E., Hagedorn, N., Manoli, G., Sekiguchi, M., Yoshii, T., Rieger, D., and Helfrich-Forster, C. (2022). The neuronal circuit of the Dorsal circadian clock neurons in *Drosophila melanogaster*. Front Physiol 13, 886432. 10.3389/fphys.2022.886432.

48. Cavanaugh, D.J., Geratowski, J.D., Wooltorton, J.R., Spaethling, J.M., Hector, C.E., Zheng, X., Johnson, E.C., Eberwine, J.H., and Sehgal, A. (2014). Identification of a circadian output circuit for rest:activity rhythms in *Drosophila*. Cell 157, 689–701. 10.1016/j.cell.2014.02.024.

49. Seluzicki, A., Flourakis, M., Kula-Eversole, E., Zhang, L., Kilman, V., and Allada, R. (2014). Dual PDF signaling pathways reset clocks via TIMELESS and acutely excite target neurons to control circadian behavior. PLOS Biology 12, e1001810. 10.1371/journal.pbio.1001810.

50. Cognigni, P., Felsenberg, J., and Waddell, S. (2018). Do the right thing: neural network mechanisms of memory formation, expression and update in *Drosophila*. Current Opinion in Neurobiology 49, 51–58. 10.1016/j.conb.2017.12.002.

51. Lyons, L.C., and Roman, G. (2009). Circadian modulation of short-term memory in *Drosophila*. Learning & memory 16, 19–27. 10.1101/lm.1146009.

52. Chouhan, N.S., Wolf, R., Helfrich-Forster, C., and Heisenberg, M. (2015). Flies remember the time of day. Current Biology: 25, 1619–1624. 10.1016/j.cub.2015.04.032.

53. Machado Almeida, P., Lago Solis, B., Stickley, L., Feidler, A., and Nagoshi, E. (2021). Neurofibromin 1 in mushroom body neurons mediates circadian wake drive through activating cAMP-PKA signaling. Nat Commun 12, 5758. 10.1038/s41467-021-26031-2.

54. Yasuyama, K., and Meinertzhagen, I.A. (2010). Synaptic connections of PDF-immunoreactive lateral neurons projecting to the dorsal protocerebrum of *Drosophila melanogaster*. J. Comparative Neurology 518, 292–304. 10.1002/cne.22210.

55. Herrero, A., Duhart, J.M., and Ceriani, M.F. (2017). Neuronal and glial clocks underlying structural remodeling of pacemaker neurons in *Drosophila*. Front Physiol 8, 918. 10.3389/fphys.2017.00918.

56. Allada, R., and Chung, B.Y. (2010). Circadian organization of behavior and physiology in *Drosophila*. Annu Rev Physiol 72, 605–624. 10.1146/annurev-physiol-021909-135815.

57. 57. Kiral, F.R., Dutta, S.B., Linneweber, G.A., Hilgert, S., Poppa, C., Duch, C., von Kleist, M., Hassan, B.A., and Hiesinger, P.R. (2021). Brain connectivity inversely scales with developmental temperature in *Drosophila*. Cell Reports 37, 110145. 10.1016/j.celrep.2021.110145.

58. Ruben, M., Drapeau, M.D., Mizrak, D., and Blau, J. (2012). A mechanism for circadian control of pacemaker neuron excitability. Journal of Biological Rhythms 27, 353–364. 10.1177/0748730412455918.

59. Groth, A.C., Fish, M., Nusse, R., and Calos, M.P. (2004). Construction of transgenic *Drosophila* by using the site-specific integrase from phage phiC31. Genetics 166, 1775–1782. 10.1093/genetics/166.4.1775.

60. Mezan, S., Feuz, J.D., Deplancke, B., and Kadener, S. (2016). PDF signaling is an integral part of the *Drosophila* circadian molecular oscillator. Cell reports 17, 708–719. 10.1016/j.celrep.2016.09.048.

61. Stuart, T., Butler, A., Hoffman, P., Hafemeister, C., Papalexi, E., Mauck, W.M., 3rd, Hao, Y., Stoeckius, M., Smibert, P., and Satija, R. (2019). Comprehensive integration of single-cell data. Cell 177, 1888–1902 e1821. 10.1016/j.cell.2019.05.031.

62. Zheng, G.X., Terry, J.M., Belgrader, P., Ryvkin, P., Bent, Z.W., Wilson, R., Ziraldo, S.B., Wheeler, T.D., McDermott, G.P., Zhu, J., et al. (2017). Massively parallel digital transcriptional profiling of single cells. Nat Commun 8, 14049. 10.1038/ncomms14049.

